# ARSDA: A new approach for storing, transmitting and analyzing high-throughput sequencing data

**DOI:** 10.1101/114470

**Authors:** Xuhua Xia

## Abstract

Two major stumbling blocks exist in high-throughput sequencing (HTS) data analysis. The first is the sheer file size typically in gigabytes when uncompressed, causing problems in storage, transmission and analysis. However, these files do not need to be so large and can be reduced without loss of information. Each HTS file, either in compressed .SRA or plain text .fastq format, contains numerous identical reads stored as separate entries. For example, among 44603541 forward reads in the SRR4011234.sra file (from a *Bacillus subtilis* transcriptomic study) deposited at NCBI’s SRA database, one read has 497027 identical copies. Instead of storing them as separate entries, one can and should store them as a single entry with the SeqID_NumCopy format (which I dub as FASTA+ format). The second is the proper allocation reads that map equally well to paralogous genes. I illustrate in detail a new method for such allocation. I have developed ARSDA software that implement these new approaches. A number of HTS files for model species are in the process of being processed and deposited at http://coevol.rdc.uottawa.ca to demonstrate that this approach not only saves a huge amount of storage space and transmission bandwidth, but also dramatically reduces time in downstream data analysis. Instead of matching the 497027 identical reads separately against the *Bacillus subtilis* genome, one only needs to match it once. ARSDA includes functions to take advantage of HTS data in the new sequence format for downstream data analysis such as gene expression characterization. ARSDA can be run on Windows, Linux and Macintosh computers and is freely available at http://dambe.bio.uottawa.ca/ARSDA/ARSDA.aspx.

## INTRODUCTION

High-throughput sequencing (HTS) is now used not only in characterizing differential gene expression, but also in many other fields where it replaces the traditional microarray approach. Ribosomal profiling, traditionally done through microarray (Arava *et al.* 2003; Mackay *et al.* 2004), is now almost exclusively done with deep sequencing of ribosome-protected segments of messages (Ingolia *et al.* 2009a; Ingolia *et al.* 2009b; Ingolia *et al.* 2011), although the results from the two approaches for ribosomal profiling are largely concordant (Xia *et al.* 2011). Similarly, EST-based (Rogers *et al.* 2012) and microarray-based (Pleiss *et al.* 2007) methods for detecting alternative splicing events and characterizing splicing efficiency is now replaced by HTS (Kawashima *et al.* 2014), especially by lariat sequencing (Awan *et al.* 2013; Stepankiw *et al.* 2015). The availability of HTS data has dramatically accelerated the test of biological hypotheses. For example, a recent study on alternative splicing (Vlasschaert *et al.* 2016) showed that skipping of exon 7 (E_7_) in human and mouse *USP4* is associated with weak signals of splice sites flanking E_7_. The researchers predicted that, in some species where the splice site signals are strong, E_7_ skipping would disappear. This prediction is readily tested and confirmed with existing HTS data, i.e., E_6_-E_8_ mRNA was found in species with weak splice signals flanking E_7_, and E_6_-E_7_-E_8_ mRNA in species with strong splice signals flanking E_7_ (Vlasschaert *et al.* 2016).

In spite of the potential of HTS data in solving practical biological problems, severe under-usage of HTS data has been reported (Team 2011). One major stumbling block in using the HTS data is the large file size. Among the 6472 transcriptomic studies on human available at NCBI/DDBJ/EBI by Apr. 14, 2016, 196 studies each contribute more than 1 Terabytes (TB) of nucleotide bases, with the top one contributing 86.4 TB. Few laboratories would be keen on downloading and analyzing this 86.4 TB of nucleotides, not to mention comparing this study to HTS data from other human transcriptomic studies.

The explosive growth of HTS data in recent years has caused serious problems in data storage, transmission and analysis(Leinonen *et al.* 2011; Kodama *et al.* 2012). Because of the high cost of maintaining such data, coupled with the fact that few researchers had been using such data, NCBI had planned the closure of the sequence read archive a few years ago (Team 2011), but continued the support only after DDBJ and EBI decided to continue their effort of archiving the data. The incident highlights the prohibitive task of storing, transmitting and analyzing HTS data, and motivated the joint effort of both industry and academics to search for data compression solutions (Janin *et al.* 2014; Zhu *et al.* 2015b; Numanagic *et al.* 2016).

## A SOLUTION WITH A NEW SEQUENCE FORMAT

HTS data files do not need to be so huge. Take for example the characterized transcriptomic data for *Escherichia coli* K12 in the file SRR1536586.sra (where SRR1536586 is the SRA sequence file ID in NCBI/DDBJ/EBI). The file contains 6,503,557 sequence reads of 50 nt each, but 195310 sequences are all identical (TGTTATCACGGGAGACACACGGCGGGTGCTAACGTCCGTCGTGAAGAGGG), all mapping exactly to sites 929-978 in *E. coli* 23S rRNA genes (The study did use the MICROBExpress Bacterial mRNA Enrichment Kit to remove the 16S and 23S rRNA, otherwise there would be many more). There are much more extreme cases. For example, one of the 12 HTS files from a transcriptomic study of *Escherichia coli* (SRR922264.sra), contains a read with 1,606,515 identical copies among its 9,690,570 forward reads. There is no sequence information lost if all these 1,606,515 identical reads are stored by a single sequence with a sequence ID such as UniqueSeqX_1606515 (i.e., SeqID_CopyNumber format which I dub as FASTA+ format with file type .fasP). Such storage scheme not only results in dramatic saving in data storage and transmission, but also leads to dramatic reduction in computation time in downstream data analysis. At present, all software packages for HTS data analysis will take the 1,606,515 identical reads separately and search them individually against the *E. coli* genome (or target gene sequences such as coding sequences). The SeqID_CopyNumber storage scheme reduces the 1,606,515 searches to a single one.

A huge chunk of SRA data stored in NCBI/DDBJ/EBI consists of ribosome profiling data (Ingolia *et al.* 2009a; Ingolia *et al.* 2009b; Ingolia *et al.* 2011), which is obtained by sequencing the mRNA segment (~30 bases) protected by the ribosome after digesting all the unprotected mRNA segments. Mapping these ribosome-protected segments to the genome allows one to know specifically where the ribosomes are located along individual mRNAs. In general, such data are essential to understand translation initiation, elongation and termination efficiencies. For example, a short poly(A) segment with about eight or nine consecutive A immediately upstream of the start codon in yeast (*Saccharomyces cerevisiae*) genes is significantly associated with ribosome density and occupancy (Xia *et al.* 2011), confirming the hypothesis that short poly(A) upstream of the start codon facilitates the recruitment of translation initiation factors but long poly(A) would bind to poly(A)-binding protein and interfere with cap-dependent translation. Sequence redundancy is high in such ribosomal profiling data and the FASTA+ format can lead to dramatic saving in the disk space of data storage and time in data transmission.

## ARSDA

I developed software ARSDA (for Analyzing RNA-Seq Data, Fig. 1a) to alleviate the problem associated with storage, transmission and analysis of HTS data. ARSDA can take input .SRA files or .fastq files of many gigabytes, build an efficient dictionary of unique sequence reads in a FASTA/FASTQ file, keep track of their copy numbers, and output them to a FASTA+ file in the SeqID_CopyNumber format (Fig. 1b). Both fixed-length and variable-length sequences can be used as input. In addition, I have implemented functions in ARSDA to take advantage of the new sequence format to perform gene expression, with the main objective to demonstrate how much faster downstream data analysis can be done with data in FASTA+/FASTQ+ format. However, ARSDA does include a unique feature in assigning shared reads among paralogous genes that I will describe below. ARSDA also includes sequence visualization functions for global base-calling quality, per-read quality and site-specific read quality (Fig. 1c-d), but these functions are also available elsewhere, e.g., FastQC (Andrews 2017) and NGSQC (Dai *et al.* 2010) and consequently will not be described further (but see the attached QuickStart.PDF).

**Fig. 1.**
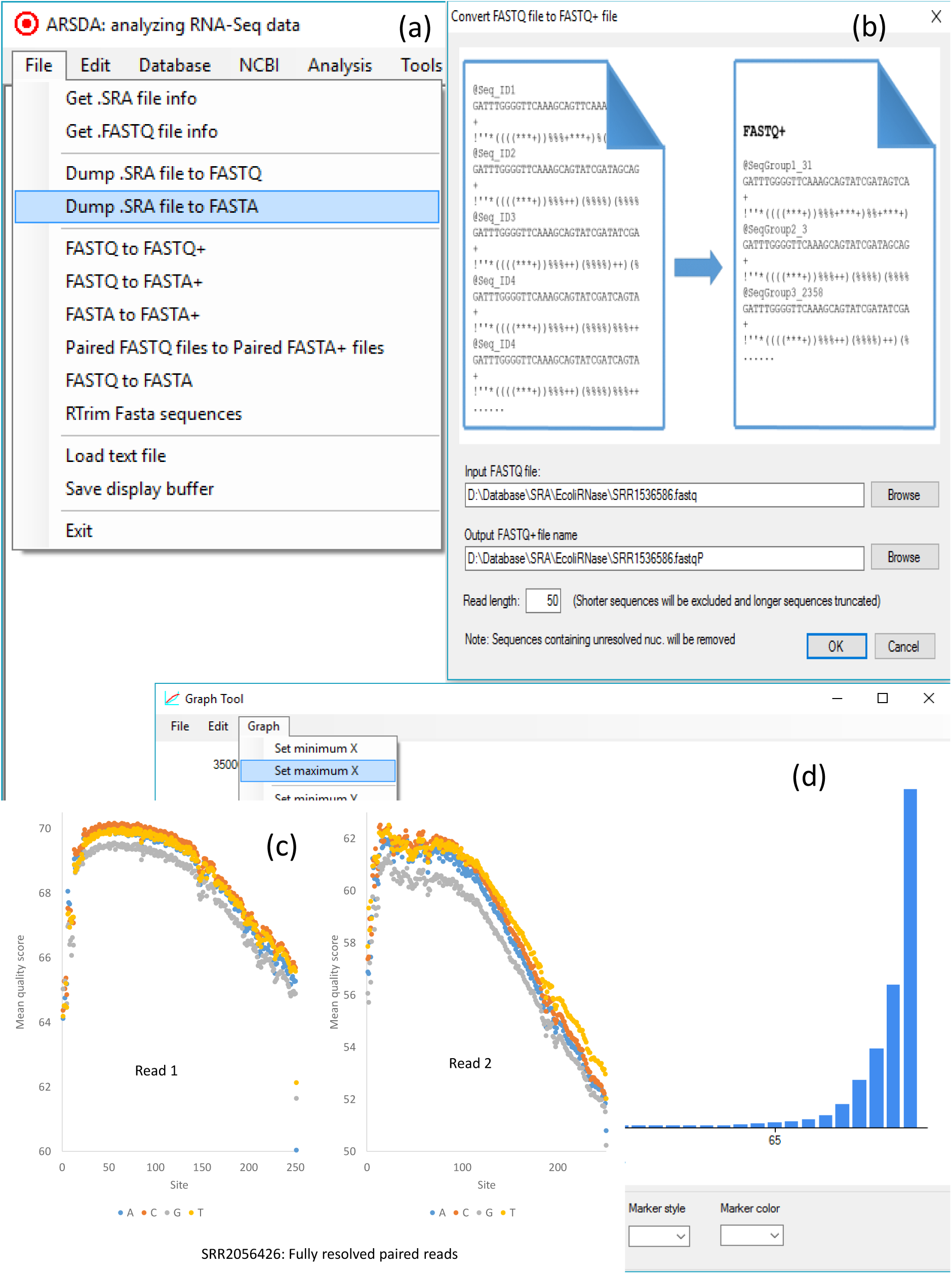
User interface in ARSDA. (a) The menu system, with database creation under the ‘Database’ menu, gene expression characterization under the ‘Analysis’ menu, etc. (b) Converting a FASTQ/FASTA file to a FASTQ+/FASTA+ file. (c) Site-specific read quality visualization. (d) Global read quality visualization.

### Converting HTS data to FASTA+/FASTQ+

The output from processing the SRR1536586.sra file (with part of the read matching output in Table 1) highlights two points. First, many sequences in the file are identical. Second, although the transcriptomic data characterized in SRR1536586 have undergone rRNA depletion by using Ambion’s MICROBExpress Bacterial mRNA Enrichment Kit (Pobre and Arraiano 2015), there are still numerous reads in the transcriptomic data that are from rRNA genes. This suggests that storing mRNA reads separately from rRNA reads can dramatically reduce file size because most researchers are not interested in the abundance of rRNAs.

**Table 1.** Part of read-matching output from ARSDA, with 195310 identical reads matching a segment of large subunit (LSU) rRNA, 86308 identical reads matching another segment of LSU rRNA, and so on. Results generated from ARSDA analysis of the SRR1536586.sra file from NCBI.

| Gene | N<sub>copy</sub> | Gene | N<sub>copy</sub> |
| --- | --- | --- | --- |
| LSU rRNA | 195310 | SSU rRNA | 30417 |
| LSU rRNA | 86308 | LSU rRNA | 29508 |
| LSU rRNA | 58400 | 5S rRNA | 28187 |
| SSU rRNA | 47323 | LSU rRNA | 24982 |
| LSU rRNA | 45695 | SSU rRNA | 23286 |
| LSU rRNA | 36258 | LSU rRNA | 19991 |
| 5S rRNA | 33674 | SSU rRNA | 19268 |

While the conversion of FASTA/FASTQ files to FASTA+ files is time-consuming, it needs to be done only once for data storage, preferably at NCBI/EBI/DDBJ, and the resulting saving in storage space, internet traffic and computation time in downstream data analysis is tremendous. The file size is 1.49 GB for the original FASTQ file derived from SRR1536586.sra, but is only 66 MB for the new FASTA+ file, both being plain text files.

I have further created BLAST databases from the processed HTS files in FASTA+ format for model species such as *E. coli, B. subtilis, S. cerevisiae* and *Caenorhabditis elegans* and deposited them at coevol.rdc.uottawa.ca. The sequence ID in these BLAST databases are in the form of SeqID_CopyNumber. These files reduce the computation time for quantifying gene expression from many hours to only a few minutes (less than two minutes for my Windows 10 PC with ani7-4770 CPU at 3.4GHz and 16 GB of RAM). This eliminates one of the key bottleneck in HTS data analysis (Liu *et al.* 2016) and would make it feasible for any laboratory to gain the power of HTS data analysis. I attach the gene expression characterized by ARSDA for the 4321 *E. coli* K12 coding sequences as supplemental file SRR1536586_GB.txt. A part of it is reproduced in Table 2.

**Table 2.**
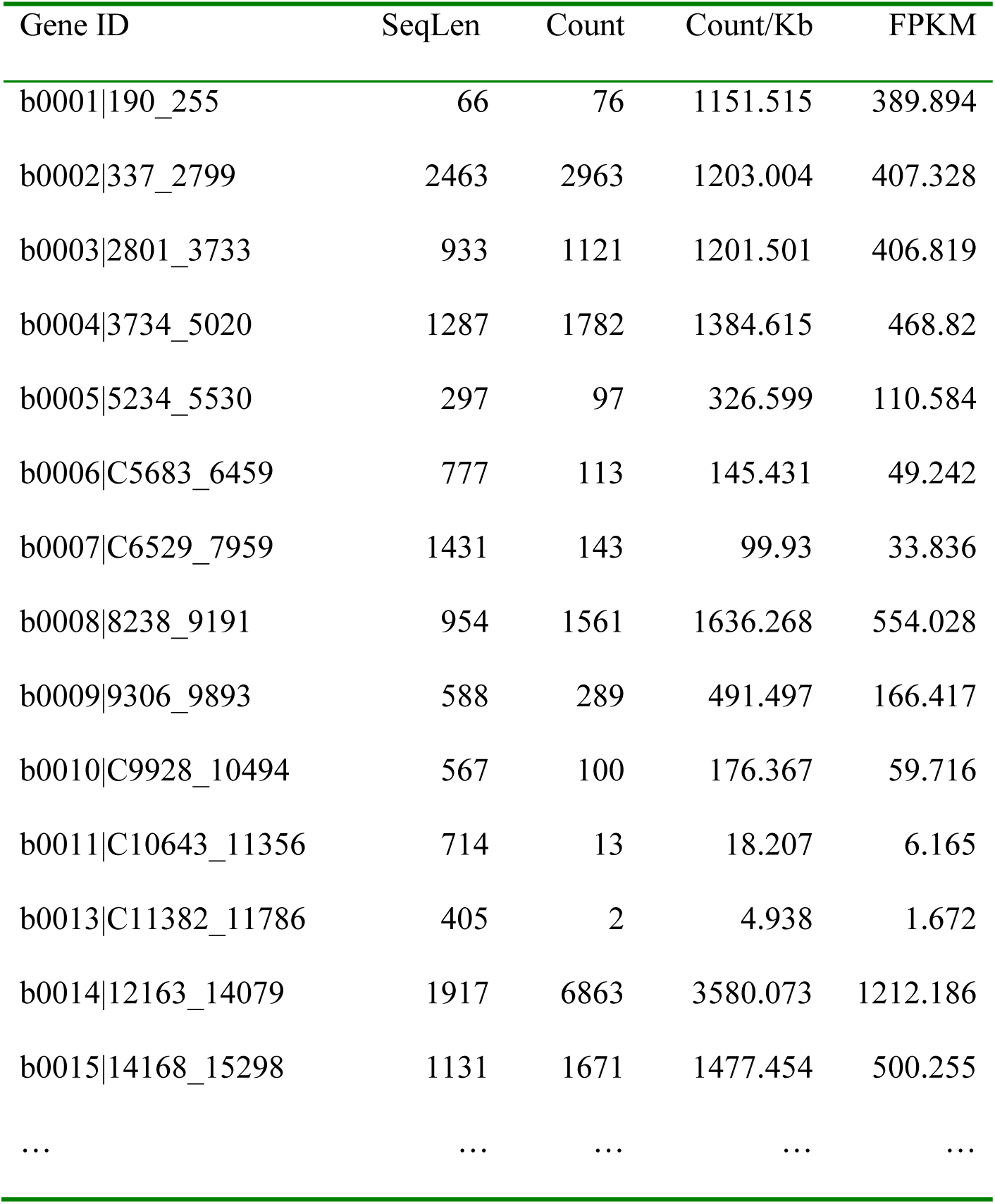
Partial output of gene expression, with the gene locus_tag (together with start and end sites) as Gene ID.

One of the frequently used sequence-compression scheme is to use a reference genome so that each read can be represented by a starting and an ending number on the genome (Benoit *et al.* 2015; Kingsford and Patro 2015; Zhu *et al.* 2015a). This approach has two problems. First, many reads do not map exactly to the genomic sequence because of either somatic mutations or sequencing errors, so representing a read by the starting and ending numbers leads to loss of information. Second, RNA-editing and processing can be so extensive that it becomes impossible to map a transcriptomic read to the genome (Abraham *et al.* 1988; Lamond 1988; Alatortsev *et al.* 2008; Li *et al.* 2009; Simpson *et al.* 2016). Furthermore, there are still many scientifically interesting species that do not have a good genomic sequence available.

### Assigning sequence reads to paralogous genes

One of the most fundamental objectives of RNA-Seq analysis is to generate an index of gene expression (FPKM: matched fragment/reads per kilobases of transcript per million mapped reads) that can be directly compared among different genes and among different experiments with different total number of matched reads (Mortazavi *et al.* 2008). The main difficulty in quantifying gene expression arises with sequence reads matching multiple paralogous genes (Trapnell *et al.* 2013; Rogozin *et al.* 2014). This problem, which has plagued microarray data analysis, is now plaguing RNA-Seq analysis. Most publications of commonly used RNA-Seq analysis methods (Langmead *et al.* 2009; Trapnell *et al.* 2009; Langmead *et al.* 2010; Roberts *et al.* 2011; Langmead and Salzberg 2012; Trapnell *et al.* 2012; Dobin *et al.* 2013; Roberts *et al.* 2013; Deng *et al.* 2014) often avoided mentioning read allocation to paralogous genes. The software tools associated with these publications share two simple options for handling matches to paralogous genes. The first is to use only uniquely matched reads, i.e., reads that match to multiple genes are simply ignored. The second is to assign such reads equally among matched genes. These options are obviously unsatisfactory. Here I describe a new approach which should substantially improve the accuracy of HTS data analysis such as gene expression characterization.

### Allocating sequence reads to paralogous genes in a two-member gene family

We need a few definitions to explain the allocation. Let L_1_ and L_2_ be the sequence length of the two paralogous genes. Let N_U.1_ and N_U.2_ be the number of reads that can be uniquely assigned to paralogous gene 1 or 2, respectively (i.e., the reads that matches one gene better than the other). Now for those reads that match the two genes equally well, the proportion of the reads contributed by paralogous gene 1 may be simply estimated as

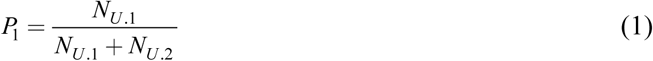

Now for any read that matches the two paralogous genes equally well, we will assign P_1_ to paralogous gene 1, and (1-P_1_) to paralogous gene 2. In the extreme case when paralogous genes are all identical, then N_u.1_ = N_u.2_ =0, and we will assign 1/2 of these equally matched read to genes 1 and 2. We should modify Eq. (1) to make it more generally applicable as follows

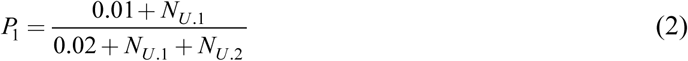

where 0.01 in the numerator and 0.02 in the denominator are pseudocounts. The treatment in Eq. (2) implies that when N_u.1_ = N_U.2_ = 0 (e.g., when two paralogous genes are perfectly identical), then a read matching equally well to these paralogous genes will be equally divided among the two paralogues.

One problem with this treatment is its assumption of L_1_ = L_2_. If paralogous gene 1 is much longer than the other, then Nm is expected to be larger than N_U.2_, everything else being equal. One may standardize N_U.1_ and N_U.2_ to number of unique matches per 1000 nt, designated by SN_U.i_ = 1000N _U.i_ /Li (where i = 1 or 2) and replace N _U.i_ in Eq. (2) by SN _U.i_ as follows (Mortazavi *et al.* 2008):

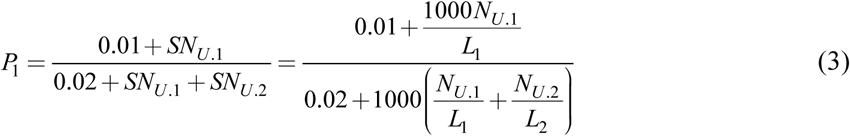

### Allocating sequence reads in gene family with more than two members

One might, mistakenly, think that it is quite simple to extend Eq. (3) for a gene family of two members to a gene family with F members by writing

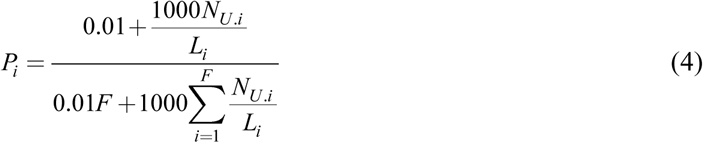

This does not work. For example, if we have three paralogous genes designated A, B, and C, respectively. Suppose that the gene duplication that gave rise to B and C occurred very recently so that B and C are identical, but A and the ancestor of B and C have diverged for a long time. In this case, N_U.B_ = N_U.C_ = 0 because a read matching B will always matches C equally well, but N_U.A_ may be greater than 0. This will result in unfair allocation of many transcripts from B and C to A according to Eq. (4). I outline the approach below for dealing with gene families with more than two members.

With three or more paralogous genes, one may benefit from a phylogenetic tree for proper allocation of sequence reads. I illustrate the simplest case with a gene family with three paralogous genes A, B, and C idealized into three segments in Fig. 3. The three genes shared one identical middle segment with 23 matched reads (that necessarily match equally well to all three paralogues). Genes B and C share an identical first segment to which 20 reads matched. Gene A has its first segment different from that of B and C and got four matched reads. The three genes also have a diverged third segment where A matched 3 reads, B matched 6 and C matched 12. Our task is then to allocate the 23 reads shared by all three and 20 reads shared by B and C to the three paralogues.

One could apply maximum likelihood or least-squares method for the estimation, but ARSDA uses a simple counting approach by applying the following

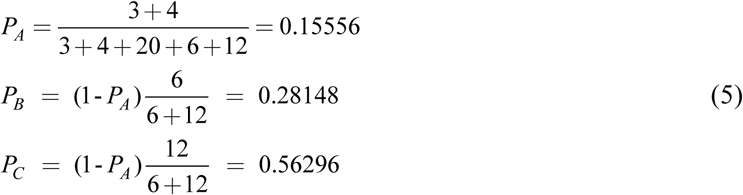

Thus, we allocate the 23 reads (that matched three genes equally) to paralogous genes A, B and C according to P_A_, P_B_ and P_C_, respectively. For the 20 reads that matched B and C equally well, we allocate 20*6/(6+12) to B and 20*12/(6+12) to C. This gives the estimated number of matches to each gene as

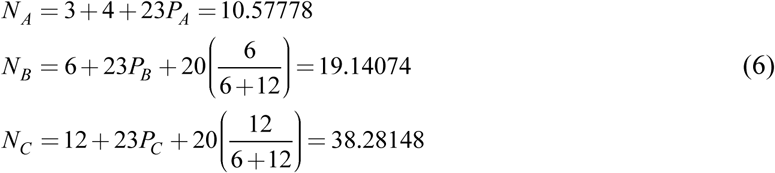

These numbers are then normalized to give FPKM (Mortazavi *et al.* 2008). The current version of ARSDA assume that gene families with more than two members to have roughly the same sequence lengths. This is generally fine with prokaryotes but may become problematic with eukaryotes.

In practice, one can obtain the same results without actually undertaking the extremely slow process of building trees for paralogous genes. One first goes through reads shared by two paralogous genes (e.g., the 20 reads shared by genes B and C in Fig. 2) and allocate the reads according to P_B_ =6/(6+12) = 1/3 and PC = 12/(6+12) = 2/3. Now genes B and C will have 12.66667 (=6+20*P_B_) and 25.33333 (=12+20*P_C_) assigned reads, i.e., N_U.B_ = 12.66667 and N_U.C_ = 25.33333. Once we have done with reads shared by two paralogous genes, we go through reads shared by three paralogous genes, e.g., the 23 reads shared by genes A, B, and C in Fig. 2. With N_U.A_ = 7, N_U.B_ = 12.66667, N_U.C_ = 25.333333, and N = N_U.A_ + N_U.B_ + N_U.C_ = 45, so we have

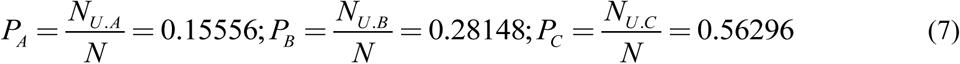

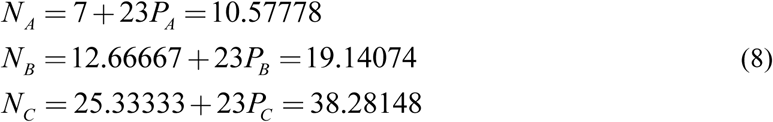

which are the same as shown in Eq. (6). This progressive process continues until we have allocated reads shared by the largest number of paralogous genes. The gene expression output in the supplemental SRR1536586_GB.txt is obtained in this way.

**Fig. 2.**
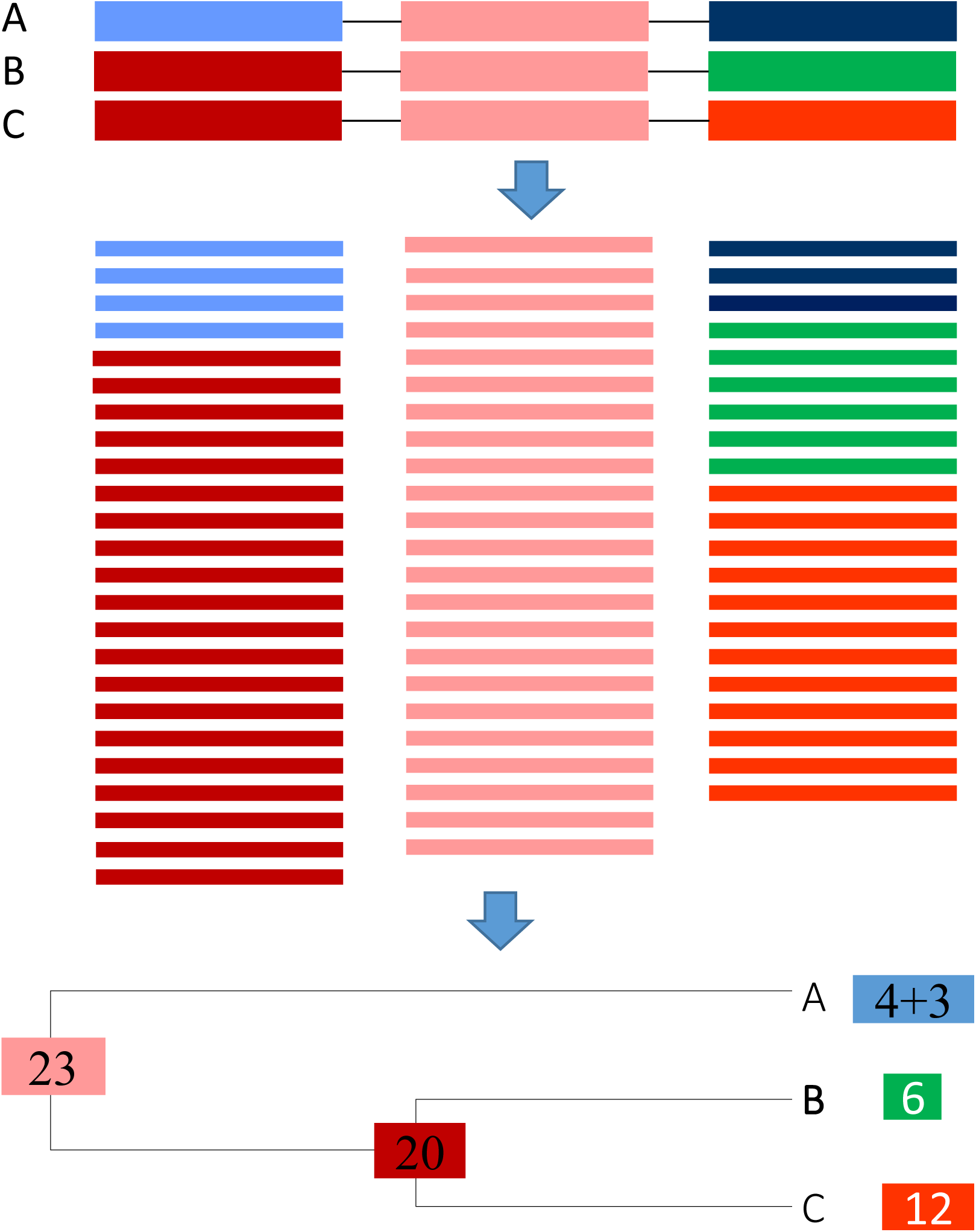
Allocation of shared reads in a gene family with three paralogous genes A, B and C with three idealized segments with a conserved identical middle segment, strongly homologous first segment that is identical in B and C, and a diverged third segment. Reads and the gene segment they match to are of the same color.

## SOFTWARE AND DATA AVAILABILITY

ARSDA is freely available at http://dambe.bio.uottawa.ca/ARSDA/ARSDA.aspx, together with a QuickStart.PDF file showing HTS file conversion from FASTA/FASTQ file to FASTA+ format, three types of HTS data quality visualization tools, and downstream characterization of gene expression. It is a Windows program but can run on any computer with .NET framework installed (e.g. Macintosh and Linux with MONO activated). The BLAST databases derived from HTS reads for several model species, in which sequence IDs are in the format of SeqID_CopyNumber, are deposited at coevol.rdc.uottawa.ca. One can use these BLAST databases with ARSDA to characterize gene expression or other analysis. Ultimately, it is NCBI/EBI/DDBJ that should store all HTS data in such BLAST databases.

## ACKNOWLEDGEMENT

This study is funded by the Discovery Grant from Natural Science and Engineering Research Council (NSERC, RGPIN/261252) of Canada. I thank J. Silke, J. Wang, Y. Wei, and C. Vlasschaert. ARSDA was demonstrated in two workshops, one organized by Dr. J. Lu of Peking University and another by Dr. E. Pranckeviciene of University of Vilnius. I thank participants for their feedback.

